# Ubiquitin-dependent recruitment of SLFN11 to chromatin is regulated by deubiquitinase (DUB) and RNF168

**DOI:** 10.64898/2026.03.26.714477

**Authors:** Daiki Taniyama, Gianluca Pegoraro, Ying Wu, Shar-Yin Naomi Huang, Craig J. Thomas, Laurent Ozbun, Andy D Tran, Liton K Saha, Junko Murai, Ukhyun Jo, Yves Pommier

**Affiliations:** Developmental Therapeutics Branch, Center for Cancer Research, National Cancer Institute, National Institute of Health, Bethesda, MD 20892, USA; High-Throughput Imaging Facility (HiTIF), Laboratory of Receptor Biology and Gene Expression, Center for Cancer Research, National Cancer Institute, National Institute of Health, Bethesda, MD 20892, USA; Advanced Biomedical Computational Science, Frederick National Laboratory for Cancer Research, Frederick, MD, 21702, USA; Division of Preclinical Innovation, Center for Cancer Research, National Cancer Institute, National Institute of Health, Bethesda, MD 20892, USA; Laboratory of Cancer Biology and Genetics, Center for Cancer Research, National Cancer Institute, National Institute of Health, Bethesda, MD 20892, USA; Department of Cell Growth and Tumor Regulation, Proteo-Science Center, Premier Institute for Advanced Studies, Ehime University, Toon, Ehime 791-0295, Japan; Pediatric Oncology Branch, Center for Cancer Research, National Cancer Institute, National Institute of Health, Bethesda, MD 20892, USA

**Keywords:** SLFN11, chromatin recruitment, ubiquitin, deubiquitinases (DUB), RNF168

## Abstract

The molecular mechanisms driving SLFN11 chromatin recruitment remain partially elucidated. Using high-throughput imaging of 162 oncology-focused compounds in U2OS cells with inducible SLFN11 expression, we discovered that deubiquitinase (DUB) inhibitors drive massive SLFN11 recruitment to chromatin, preferentially at promoter regions while concurrently suppressing transcription. DUB inhibitors such as VLX-1570 promote ubiquitin-dependent enrichment of SLFN11 without detectable DNA damage, distinct from the camptothecin-induced RPA-associated SLFN11 foci formed at stressed replication forks. Yet, SLFN11 chromatin recruitment both by DUB inhibitors and DNA damage are suppressed by TAK243 demonstrating their ubiquitylation dependency. RNF168 is required for SLFN11 ubiquitylation and its subsequent chromatin association, and ubiquitylation within SLFN11’s middle linker domain (lysines 390, 391, and 429) with K27-linked polyubiquitin chains is essential for the chromatin recruitment of SLFN11. These findings suggest the importance of SLFN11 ubiquitylation by RNF168 for SLFN11 chromatin recruitment and SLFN11 transcriptional regulatory role at promoter regions.

## INTRODUCTION

Schlafen 11 (SLFN11) is a critical factor determining the vulnerability of cells to various DNA-damaging agents (DDAs) producing replication stress: inhibitors of topoisomerase I (TOP1) [camptothecin (CPT) and its clinical derivatives], of TOP2 (etoposide), of poly(ADPribose) polymerase (PARP) (olaparib, niraparib, talazoparib, rucaparib), ribonucleotide reductase (gemcitabine) as well as antimetabolites (hydroxyurea) and DNA crosslinkers agents (platinum derivatives) ^1–3^. Retrospective analyses of clinical samples have further demonstrated that high SLFN11 expression is associated with improved responses to DDAs in multiple cancer types ^4–11^.

Upon exposure to DDAs, SLFN11 is recruited to chromatin at sites of DNA damage in association with replication protein A (RPA) filaments, triggering an irreversible and lethal replication block ^12–14^. By contrast, SLFN11-deficient cells are overall resistant to DDAs while remaining sensitive to tubulin and protein kinase inhibitors ^15^. The recruitment of SLFN11 to chromatin under replication stress induces the degradation of the replication initiation factor CDT1 ^16^, inhibits ATR synthesis via tRNA cleavage ^17^, enhances chromatin accessibility ^18^, activates the early stress response including JUN, FOS and p21 ^18^, drives the degradation of reversed replication forks ^19^ and alleviates proteotoxic stress ^16^. Additionally, SLFN11 induces TP53-independent apoptosis by impairing ribosome biogenesis and inhibiting translation during the DNA damage response (DDR) ^20,21^. SLFN11 have been found localize not only to stressed replication forks but also to non-replicating chromatin regions, including promoters, intergenic regions, and untranslated regions ^22^. However, the underlying molecular mechanisms regulating its chromatin recruitment remain unclear. Therefore, elucidating these pathways is important to understand SLFN11’s role in genome regulation and cellular response mechanisms.

Post-translational modifications (PTMs) are key regulators of protein functions, stability and interactions. They include phosphorylation, methylation, acetylation, glycosylation, and ubiquitylation, each impacting cellular processes in different ways. Dephosphorylation of SLFN11 has been reported to regulate its activity ^23^, and structural analyses have shown that phosphorylation at site S753 acts as a conformational switch controlling SLFN11 dimerization and interactions with ATP and single-stranded DNA (ssDNA) ^24^. SLFN11 phosphorylation at residues S219 and T230 also controls tRNA recognition and nuclease activity.

Ubiquitylation is also a critical post-translational modification regulating protein stability and interactions through the attachment of mono- or poly-ubiquitin chains ^25^. Ubiquitylation is a multi-step process orchestrated by E1 ubiquitin-activating enzymes, E2 ubiquitin-conjugating enzymes and an E3 ubiquitin ligases. Ubiquitin molecules form poly-ubiquitin chains through various lysine linkages (M1, K6, K11, K27, K29, K33, K48, and K63), which determine their function. Lys63-linkages drive protein-protein interactions, and Lys11-, Lys29-, and Lys48-linkages target proteins for proteasomal degradation ^26^. K27-linked ubiquitylation, together with K63-linked chains regulates chromatin remodeling, as well as protein interaction, translocation and activation ^27,28^. Ubiquitylation is a reversible and dynamic process regulated by deubiquitinases (DUBs)—a diverse family of approximately 100 proteases that remove ubiquitin moieties from substrates, ensuring the precise regulation of cellular functions and protein homeostasis.

SLFN11 ubiquitylation has remained largely unexplored. The Human cytomegalovirus (HCMV) protein RL1 has been shown to induce SLFN11 degradation as part of a viral strategy to evade antiviral innate immunity ^29^. As a viral DCAF (DDB1- and CUL4-Associated Factor) HCMV RL1 recruits the CRL4 E3 ligase complex to mediate SLFN11 ubiquitylation and proteasomal degradation. However, the functional implications of SLFN11 ubiquitylation, particularly in relation to its roles in chromatin recruitment, are unknown.

In this study, we employed a high throughput pharmacological screening approach to discover novel factors driving the recruitment of SLFN11 to chromatin. After generating doxycycline-inducible SLFN11-expressing U2OS cells, which normally do not express SLFN11, we performed a spinning-disk microscopy screen ^30^ and found that DUB inhibitors are the most potent inducers of SLFN11 recruitment to chromatin (more effective than CPT, Prexasertib or doxorubicin), particularly at promoter regions. Mechanistic studies show that SLFN11 chromatin recruitment requires its ubiquitylation through K27 linkage mediated by RNF168 as an E3 ligase targeting several lysine residues in SLFN11’s middle domain. These findings reveal the importance of ubiquitin-mediated post-translational modifications in regulating the chromatin recruitment of SLFN11.

## RESULTS

### High-Throughput Imaging (HTI) microscopy reveals that deubiquitinase inhibitors promote extensive chromatin recruitment of SLFN11

High-throughput imaging (HTI) microscopy-based screen was performed with 162 oncology-focused small molecules targeting epigenetic regulation, post-translational modifications and transcription factor modulation, as well as positive control drugs such as CPT known to induce replicative stress and SLFN11 foci ^1,15^. Additional consideration was given to the medical regulatory status of the drugs, with priority assigned to approved drugs or agents currently in advanced clinical trials (**Table S1**). After doxycycline induction (5 µg/mL for 72 hours) (**Fig. 1A**), drugs were added individually to 384-well plates at three concentrations (0.1, 1 and 10 µM) for 2 hours. A pre-extraction step was included to only monitor chromatin-associated SLFN11 ^13–15^. Chromatin-bound SLFN11 was quantified using automated microscopy, and the intensity ratios were calculated by comparing the mean intensity of drug-treated samples to untreated controls (**Fig. 1B**).

**Fig. 1.**
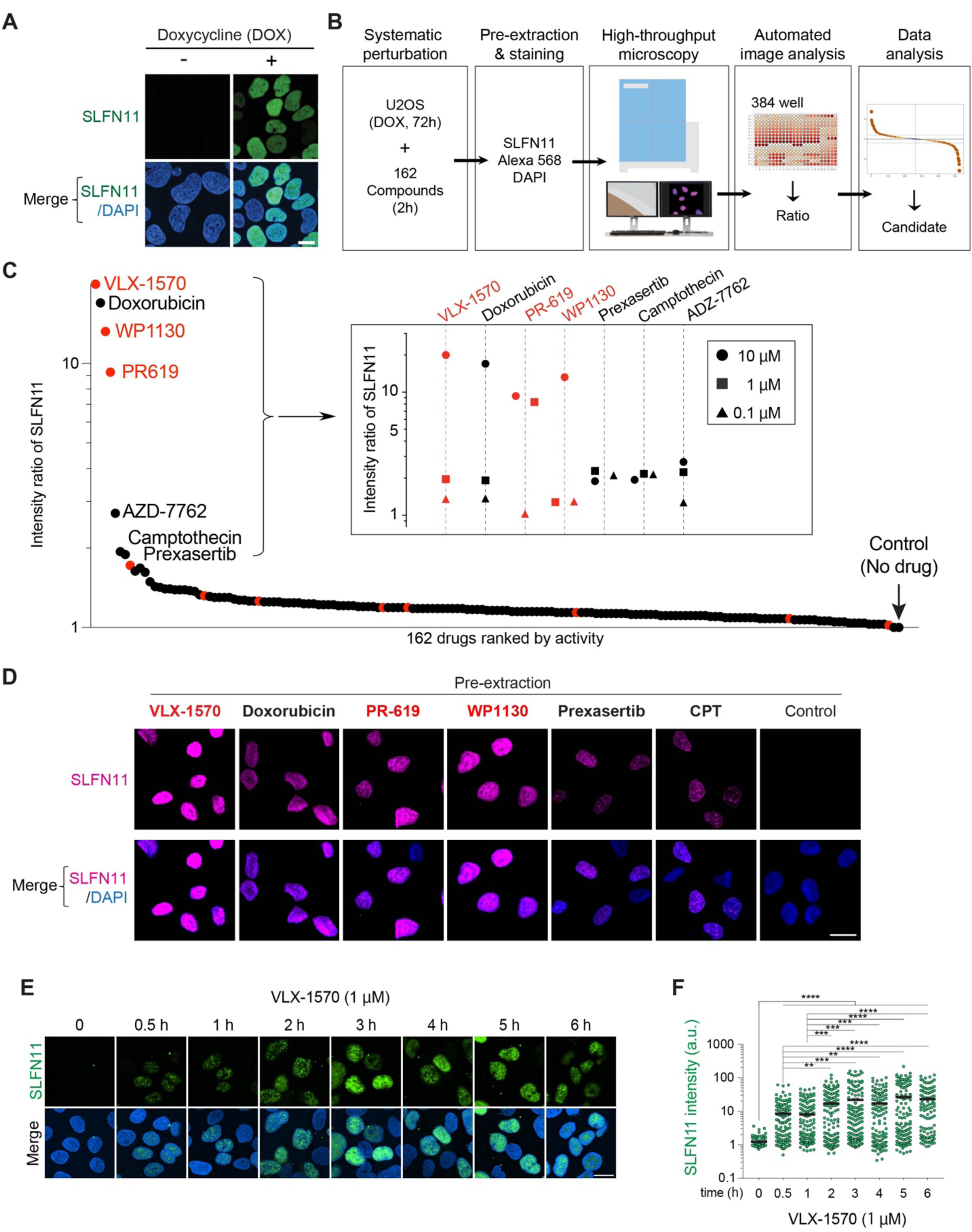
High-throughput imaging assay for SLFN11 chromatin recruitment. SLFN11 chromatin binding is induced by different DUB inhibitors. (A) Representative immunofluorescence images in doxycycline (DOX)-inducible SLFN11-expressing U2OS cells (5 µg/ml DOX, 72 hrs). Without pre-extraction. (B) Schematic overview of the drug screen workflow. (C) Scatter plot showing the drug screen results. Intensity ratio of SLFN11 at 10 µM concentration are shown. For top ranked 7 drugs, intensity ratio of three concentrations are shown (0.1, 1 and 10 µM; see inset). At minimum, 950 cells were analyzed for each condition. (D) Representative immunofluorescence images from the drug screen. Cells were washed with pre-extraction buffer before fixation to only detect chromatin-bound SLFN11 by confocal microscopy (magenta). CPT (camptothecin) and Prexasertib were used as positive controls. Scale bar: 20 µm. (E) Representative immunofluorescence images of a VLX-1570 time-course experiment. Scale bar: 10 µm. DOX-inducible SLFN11-expressing U2OS cells were treated with VLX-1570 (1 µM) for the indicated times. (F) Quantification of chromatin-bound SLFN11 signals in individual cells for the indicated treatments depicted in panel E (n = 115 – 154 cells per condition). **p < 0.01, ***p < 0.001, ****p < 0.0001 (one-way ANOVA). a.u., arbitrary units.

The HTI screen identified three deubiquitinase (DUB) inhibitors (VLX-1570, PR-619 and WP1130) that produced extensive SLFN11 chromatin recruitment (**Fig. 1C**). Notably, the DUB inhibitors were markedly more potent than CPT, VLX-1570, PR-619, and WP1130 displayed a pan-nuclear staining pattern, which contrasted with the known SFLN11 foci patterns induced by DDAs (**Fig. 1D and S1A**) ^14,18^. These observations suggested that upon DUB inhibition, SLFN11 is recruited across chromatin beyond stressed DNA replication forks. Confocal microscopy validation of the screening results confirmed that the top DUB inhibitors, including the USP14/UCHL5 inhibitor VLX-1570, the USP7/USP21 inhibitor BAY11-7082 and the broader selectivity DUB inhibitors, PR-619 and WP1130 markedly enhanced SLFN11 recruitment to chromatin (**Fig. S1B, C**). In contrast, DUB inhibitors targeting individual enzymes (UCHL1, USP1 and USP7) showed weaker or no effect, suggesting that simultaneous inhibition of multiple DUBs contribute to SLFN11 chromatin recruitment. As expected, we confirmed that, under these conditions, the DUB inhibitors led to accumulation of ubiquitylated proteins in both whole-cell and chromatin-fractionated lysates (**Fig. S1D**).

Notably, kinetics of SLFN11 chromatin recruitment in response to VLX-1570 was initiated within 30 minutes of VLX-1570 treatment (**Fig. 1E, F**), considerably faster than in response to DDAs, which typically require over 2 hours ^13,14^. We also observed that SLFN11 chromatin recruitment persisted for up to 24 hours after removal of VLX-1570 but was reversed by 96 hours (**Fig. S1E-G**). Together, these results indicate that DUB inhibition triggers the rapid yet reversible SLFN11 chromatin recruitment, in contrast to the slower and more persistent SLFN11 chromatin recruitment induced by DDAs.

### The sites of SLFN11 chromatin recruitment induced by DUB inhibitor differ from those induced by replication stress

Given that DUB inhibitors generate distinct nuclear microscopic patterns from DDAs (see Fig. 1 and S1), we hypothesized that this recruitment occurs at sites spatially distinct from the SLFN11 foci previously observed in response to replication stress agents such as CPT ^14,18^, which depend on the RPA complex binding to single-stranded DNA ^13,14^. To test this, we examined whether DUB inhibitor–induced SLFN11 chromatin recruitment is associated with RPA foci. In DU145 prostate cancer cells, which endogenously express high levels of SLFN11, and where SLFN11 largely co-localize with RPA2 foci in response to CPT, only a small fraction of the VLX-1570–induced SLFN11 chromatin foci overlapped with RPA2 (**Fig. 2A**), a pattern that was recapitulated in doxycycline-inducible U2OS cells (**Fig. 2B**). Although VLX-1570 treatment increased RPA2 chromatin foci, unlike CPT, we observed no significant association between SLFN11 and RPA2 (**Fig. S2**). Together, these findings indicate that SLFN11 chromatin recruitment by DUB inhibitors occurs largely independently of RPA foci and replication forks.

**Fig. 2.**
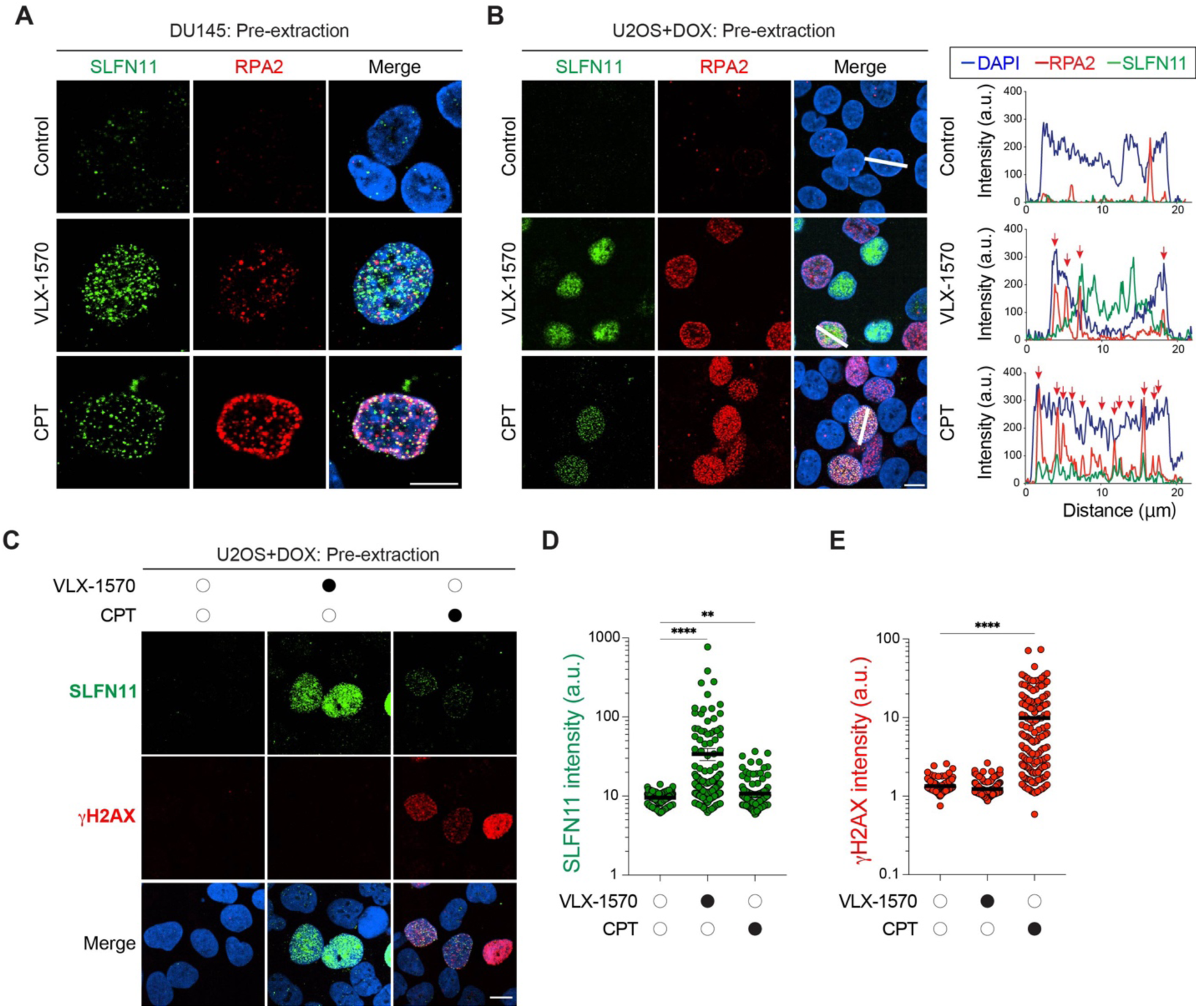
SLFN11 chromatin recruitment by DNA damage is dependent on both ubiquitin and RPA. (A) SLFN11 chromatin recruitment by VLX-1570 and camptothecin (CPT). Representative immunofluorescence images show chromatin-bound SLFN11 (green), RPA2 (red), and DAPI (blue). DU145 cells were treated with VLX-1570 (1 µM) or CPT (10 µM) for 4 hrs. Scale bar: 10 µm. (B) VLX-1570 recruits SLFN11 in an RPA-independent manner. Left: Representative immunofluorescence images showing chromatin-bound SLFN11 (green), RPA2 (red), and DAPI (blue). Scale bar: 10 µm. DOX-induced SLFN11-expressing U2OS cells were treated with VLX-1570 (1 µM) or CPT (10 µM) for 4 hrs. Right: Tracing of the distribution of signals along the white dashed lines shown in the left panels. (C) Representative immunofluorescence images showing chromatin-bound SLFN11 (green), γH2AX (red) and DAPI (blue). DOX-inducible SLFN11-expressing U2OS cells were treated with VLX-1570 (1 µM) or CPT (10 µM) for 4 hours. Scale bar: 10 µm. (D, E) Quantification of chromatin-bound SLFN11 and γH2AX signals in individual cells for the indicated treatments. (n = 118 – 166 cells per condition) *p < 0.05, **p < 0.01 **p < 0.001. ****p < 0.0001 (two-tailed unpaired t test). a.u., arbitrary units.

This conclusion is further supported by the differential phosphorylation of histone H2AX, a canonical DNA damage response marker ^31^. Whereas CPT treatment induces both SLFN11 and γH2AX foci ^13,14^, VLX-1570 elicits robust SLFN11 chromatin recruitment without detectable γH2AX signal (**Fig. 2C-E**), implying that DUB inhibitor-induced SLFN11 chromatin recruitment occurs independently of DNA damage signaling.

### VLX-1570 promotes SLFN11 recruitment associated with H3K27ac-defined open chromatin and transcription suppression

To determine the chromatin regions associated with DUB inhibitor–induced SLFN11 recruitment, we co-stained cells for SLFN11 and H3K27ac (histone H3K27 acetylation), a marker of active transcription and open chromatin. VLX-1570–treated cells showed enhanced H3K27ac chromatin staining together with SLFN11 chromatin recruitment (**Fig. 3A)**. By contrast, H3K27me3, a marker of heterochromatin, remained unchanged and showed no association with SLFN11 upon VLX-1570 treatment (**Fig. S3A, B**). These results suggest that the SLFN11 chromatin recruitment induced by VLX-1570 is associated with chromatin opening.

**Fig. 3.**
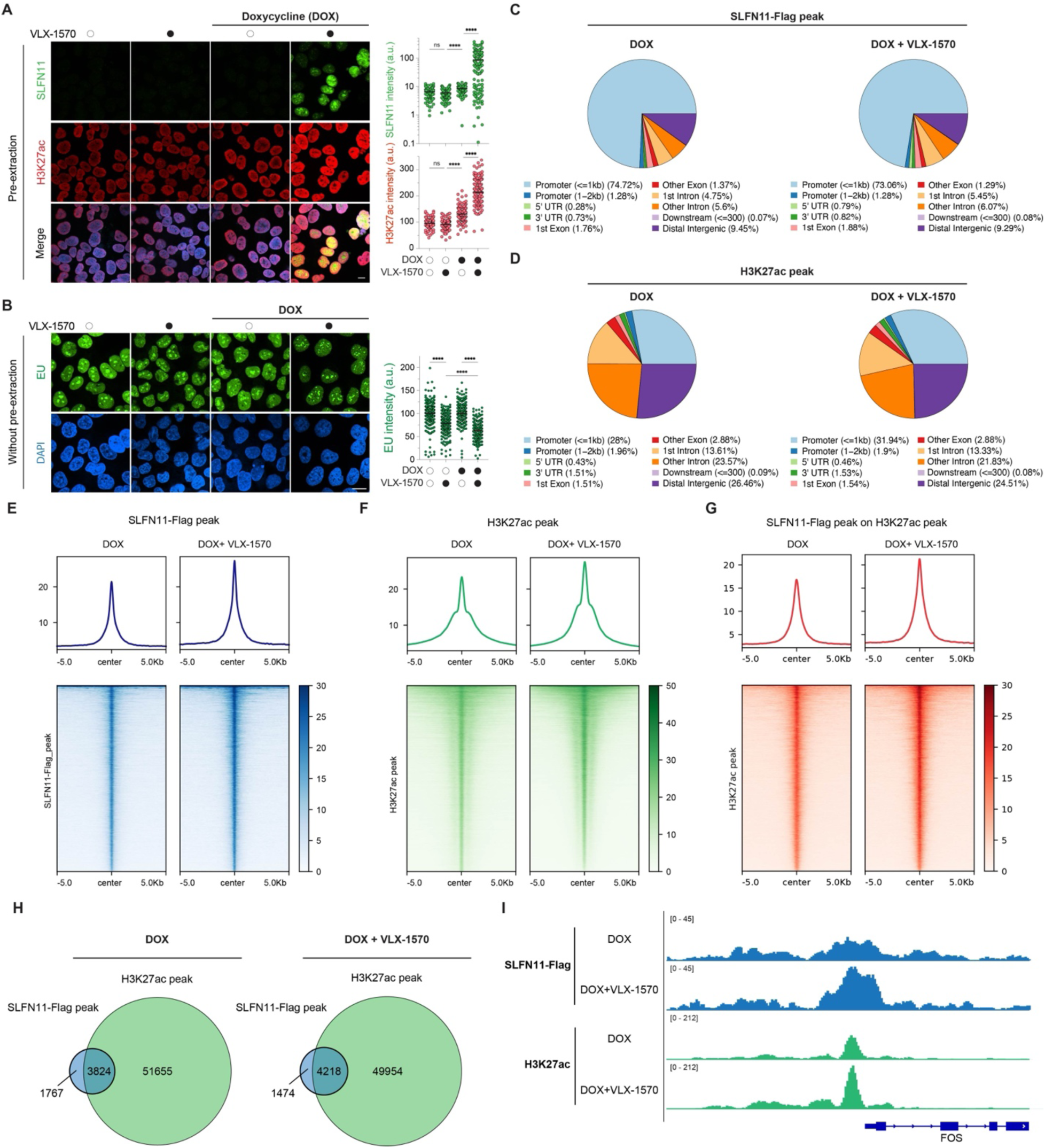
SLFN11 chromatin binding induced VLX-1570 localizes in open chromatin at promoters and suppresses RNA synthesis. (A) Left: Representative immunofluorescence images showing the colocalization of chromatin-bound SLFN11 (green) and H3K27ac (red). Nuclei are stained with DAPI (blue). DOX-induced SLFN11-expressing U2OS cells were treated with VLX-1570 (1 µM) for 4 hrs. Right: Quantification of chromatin-bound SLFN11 and H3K27ac in individual cells for the indicated treatments (n = 228 – 327 cells per condition). **p < 0.01, ns, non-significant, ***p < 0.001, ****p < 0.0001 (one-way ANOVA). a.u., arbitrary units (B) Left: Representative immunofluorescence images showing SLFN11-dependent inhibition of EU incorporation (green) by VLX-1570. Scale bar: 20 µm. DOX-inducible SLFN11-expressing U2OS cells were treated with VLX-1570 (1 µM). Nuclei are stained with DAPI (blue). Right: Quantification of EU in individual cells. (n = 109 – 166 cells per condition). **p < 0.01, ***p < 0.001, ****p < 0.0001 (one-way ANOVA) a.u.., arbitrary units (C, D) Pie charts depicting the genomic distribution of SLFN11-Flag (C) and H3K27ac (D) ChIP-seq peaks across annotated genomic features in the presence or absence of VLX-1570. (E, F, and G) Heatmaps and average signal profiles of SLFN11-Flag ChIP-seq (E), H3K27ac ChIP-seq (F), and SLFN11-Flag signal centered on H3K27ac peaks (G) with or without VLX-1570 treatment. (H) Venn diagram showing the number of SLFN11-Flag peaks, H3K27ac peaks, and their overlap identified in the indicated ChIP-seq analyses. 68.4% and 74.1% of the SLFN11 peaks coincide with the H3K27 peaks in the absence and presence of VLX-1570, respectively. (I) Genome browser tracks illustrating ChIP-seq signal enrichment at the FOS promoter for SLFN11-Flag and H3K27ac in the presence or absence of VLX-1570.

To assess the functional consequences of SLFN11 recruitment with open chromatin upon DUB inhibition, we examined nascent RNA synthesis using 5-ethyl uridine (EU) pulse-labeling and click chemistry as a readout of transcriptional activity. VLX-1570 markedly reduced nascent RNA synthesis both in the absence and presence of SLFN11, displaying a more pronounced effect upon chromatin recruitment of SLFN11 (**Fig. 3B)**. These results suggest that VLX-1570-induced SLFN11 chromatin recruitment may function as a negative regulator of transcription, similar to its established role in response to replication stress.

To map the recruitment of SLFN11 with chromatin, we performed ChIP-seq in DOX-induced SLFN11-expressing cells with or without VLX-1570 treatment. SLFN11 peaks were predominantly localized at promoter regions, whereas H3K27ac exhibited a broader distribution consistent with its presence at multiple regulatory elements, including enhancers and super-enhancers (**Fig. 3C, D**). VLX-1570 treatment markedly increased SLFN11 chromatin occupancy as well as global H3K27ac levels, with strong enrichment of SLFN11 at H3K27ac-marked regions, where approximately 80% of SLFN11 peaks overlapped with H3K27ac peaks (**Fig. 3E–H**). This co-occupancy is exemplified at the FOS promoter, a known transcriptionally responsive locus linked to SLFN11 function ^18^ (**Fig. 3I**). Collectively, these results demonstrate that SLFN11 is preferentially recruited to transcriptionally active promoters within open chromatin in response to DUB inhibition by VLX-1570.

### Ubiquitylation is required for SLFN11 recruitment to chromatin

Given the central role of DUB enzymes in ubiquitin signaling, we next examined whether SLFN11 chromatin recruitment depends on ubiquitylation. Using TAK-243, a selective inhibitor of the ubiquitin-activating enzyme UBA1 ^32^, we observed that inhibition of ubiquitylation abrogated SLFN11 recruitment induced by both CPT and the pan-DUB inhibitor PR619 (**Fig. 4A, B**). These results show that ubiquitylation is critical for SLFN11 recruitment to chromatin.

**Fig. 4.**
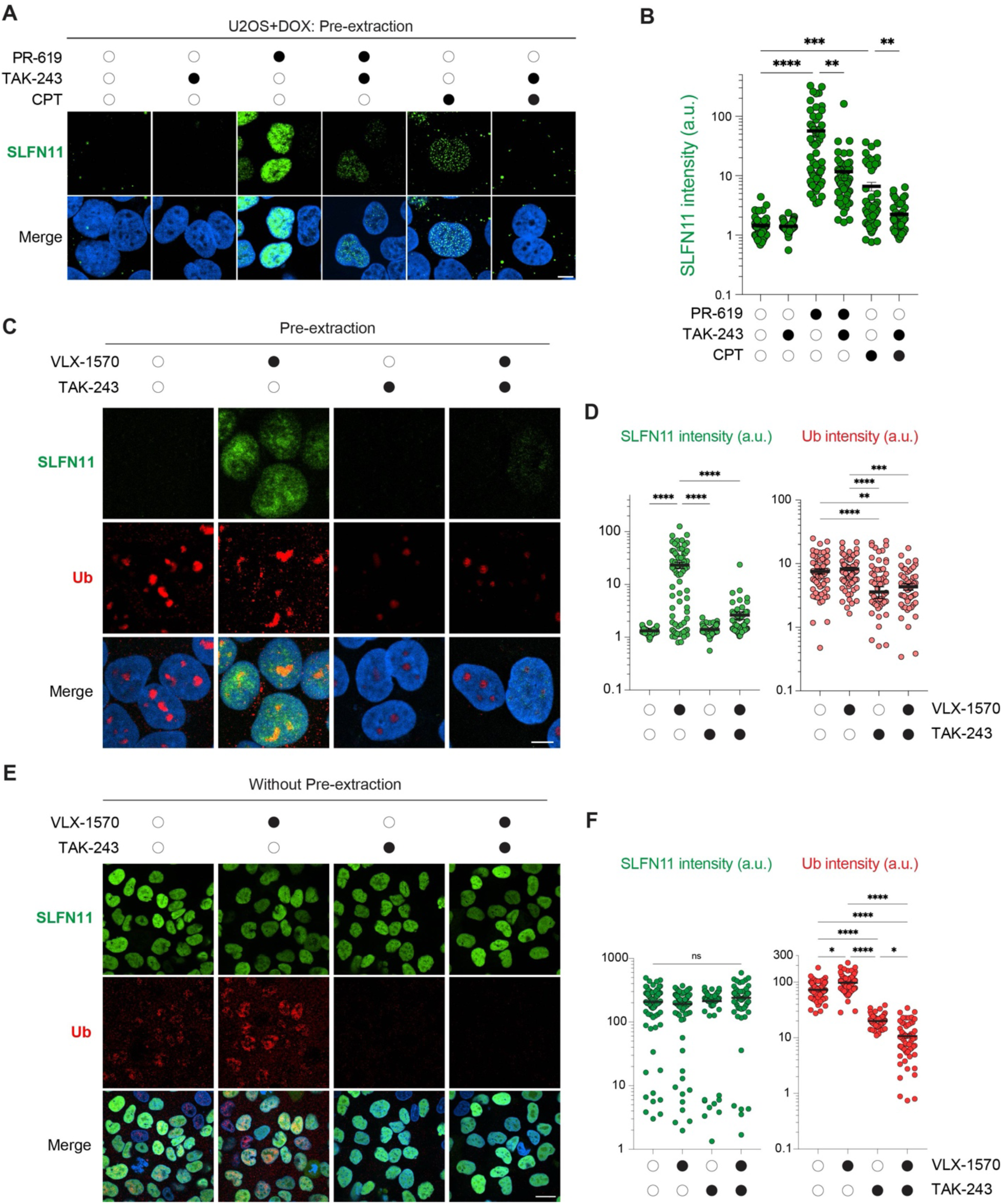
SLFN11 chromatin binding by DUB inhibitor is ubiquitination dependent. (A) Representative immunofluorescence images showing chromatin-bound SLFN11 (green) and DAPI (blue). DOX-inducible SLFN11-expressing U2OS cells were treated with PR-619 (10 µM, 4 hrs), CPT (10 µM, 4 hrs), and TAK243 (2 µM, 5 hrs). Scale bar: 5 µm. (B) Quantification of chromatin-bound SLFN11 signals in individual cells for the indicated treatments depicted in panel A (n = 38 – 77 cells per condition). **p < 0.01, ***p < 0.001, ****p < 0.0001 (two-tailed unpaired t test). a.u., arbitrary units. (C) Representative immunofluorescence images showing chromatin-bound SLFN11 (green), Ub (red) and DAPI (blue). DOX-inducible SLFN11-expressing U2OS cells were treated with VLX-1570 (1 µM, 4 hrs) with or without 1 hr pre-treatment with TAK243 (2 µM). Scale bar: 5 µm. (D) Quantification of chromatin-bound SLFN11 and Ub signals in individual cells for the indicated treatments depicted in panel C (n = 58 – 77 cells per condition). ****p < 0.0001 (two-tailed unpaired t test). a.u., arbitrary units. (E) Representative immunofluorescence images showing unchanged total SLFN11 levels (without pre-extraction; green), Ub (red) and DAPI (blue). DOX-inducible SLFN11-expressing U2OS cells were treated with same condition as shown in panel C. Scale bar: 10 µm. (F) Quantification of total SLFN11 and Ub signals in individual cells for the indicated treatments depicted in panel E (n = 60 – 81 cells per condition). *p < 0.05, ****p < 0.0001 (two-tailed unpaired t test). a.u., arbitrary units.

Under the conditions where pretreatment with TAK-243 significantly impaired SLFN11 recruitment to chromatin, the cellular ubiquitin immunofluorescence staining was reduced (**Fig. 4C, and S4A, B),** suggesting that pre-existing ubiquitin-dependent processes are essential for chromatin opening and subsequent SLFN11 loading. In contrast, total cellular SLFN11 levels, assessed without chromatin pre-extraction, remained unchanged under VLX-1570, TAK-243 or combined treatment conditions (**Fig. 4E, F**), indicating that ubiquitin signaling specifically regulates chromatin recruitment without significantly affecting the overall SLFN11 abundance. Collectively, these findings demonstrate that the ubiquitylation is required for DUB inhibitor-induced SLFN11 recruitment to chromatin.

### Ubiquitin lysine K27 is a key determinant of SLFN11 ubiquitylation

Following the observation that VLX-1570 induces chromatin opening and promotes SLFN11 recruitment, we directly tested SLFN11 ubiquitylation. Accordingly, immunoblotting revealed accumulation of high molecular-weight SLFN11 bands in chromatin fractions upon VLX-1570 treatment (**Fig. S5A**).

Next, we performed Ni-NTA–based ubiquitin pull-down assays following transient expression of His-tagged ubiquitin. SLFN11 ubiquitylation was readily detected and further enhanced by treatment with MG132 (**Fig. 5A**), indicating that SLFN11 this approach could be used to determine the specific ubiquitin linkage types involved. Because polyubiquitin chains can be assembled through multiple lysine residues (K6, K11, K27, K29, K33, K48, and K63) (**Fig. 5B**) ^33^, we generated a series of His-ubiquitin lysine-to-arginine mutants to disrupt specific linkage types. Abrogating the K27 linkages with the K27 (K27R) mutated construct markedly reduced SLFN11 ubiquitylation, whereas the K11R or K48R mutations led to increased SLFN11 ubiquitination (**Fig. 5C, D and S5B**). These findings suggest that K27-linked ubiquitylation may precede or facilitate subsequent ubiquitin modifications, while the canonical K48 and K11-linked chains contribute to proteasomal recognition and turnover.

**Fig. 5.**
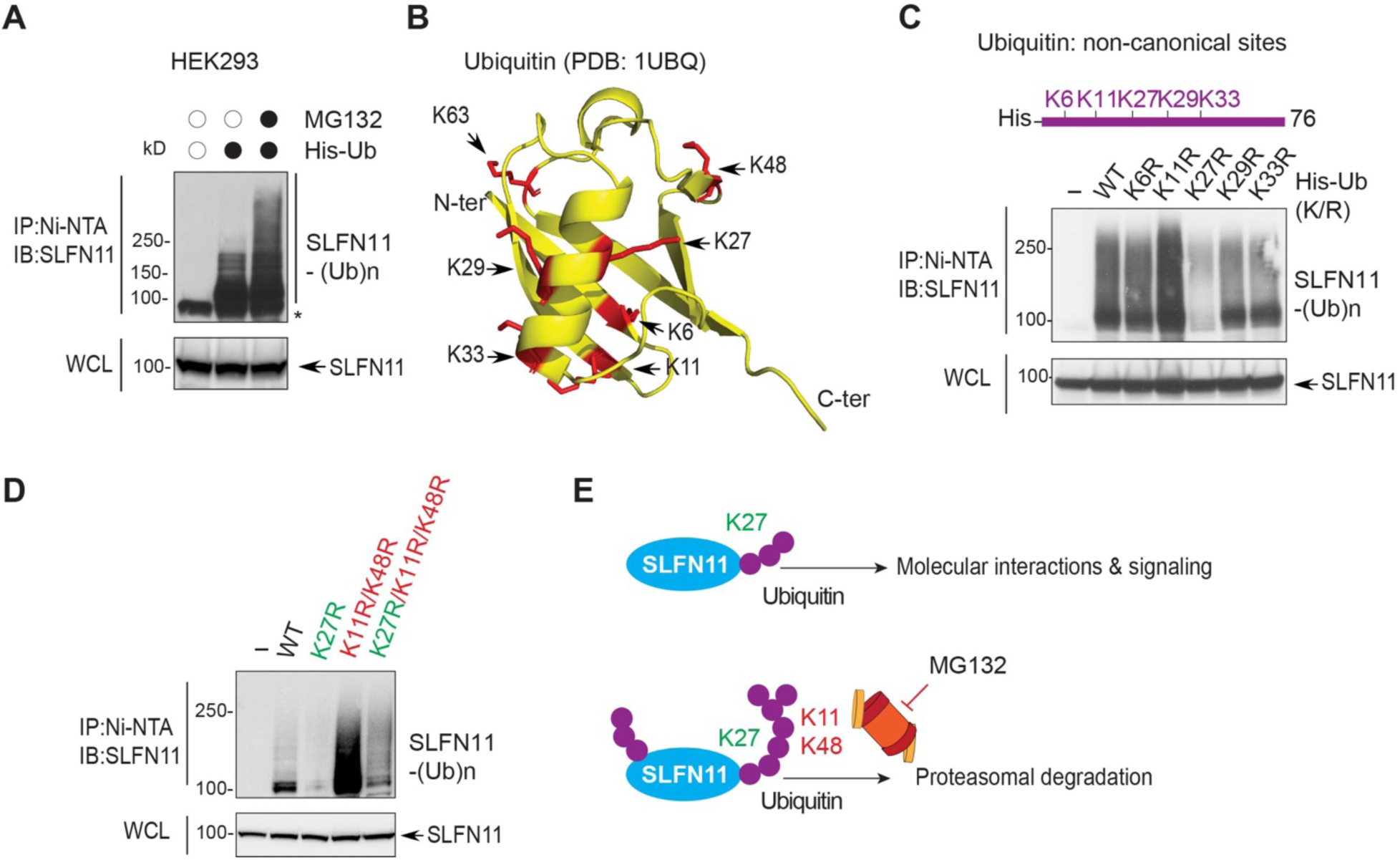
SLFN11 is ubiquitylated via K27, K11, and K48 ubiquitin linkages. (A) Cellular ubiquitylation of SLFN11. HEK293 cells were transfected with His-ubiquitin (His-Ub) and subsequently treated with or without the proteasome inhibitor MG132 (20 µM, 4 hrs). Cells were lysed and immunoprecipitated with Ni-NTA beads. Ubiquitylated SLFN11 [SLFN11-(Ub)n] was detected with anti-SLFN11 antibody. (B) Structure of ubiquitin and mapping of the 7 lysine residues (Lys6, Lys11, Lys27, Lys29, Lys33, Lys48, and Lys63) involved in protein ubiquitylation (PDB: 1UBQ). (C) Ubiquitylation of SLFN11 depends on the non-canonical ubiquitin K27 linkage. HEK293 cells were transfected with either His-Ub or with the non-canonical His-Ub K/R mutants (K6R, K11R, K27R, K29R, and K33R). Cells were lysed and immunoprecipitated with Ni-NTA beads. Ubiquitylated SLFN11 was detected with anti-SLFN11 antibody. (D) K27 ubiquitin linkages are critical for SLFN11 ubiquitylation. HEK293 cells were transfected with either His-Ub or His-Ub K/R mutants (K27R, K11R/K48R, and K27R/K11R/K48R). Cells were lysed and immunoprecipitated with Ni-NTA beads. SLFN11-(Ub)n was detected with anti-SLFN11 antibody. (E) Proposed schematic diagram of the subsequential ubiquitin linkages leading to SLFN11 chromatin recruitment (top) or proteasomal degradation (bottom).

Although K48- and K63-linked ubiquitin chains are well-established mediators of proteasomal degradation, the functional role of non-canonical K27 linkages remains less clear. To further dissect this, we generated combinatorial ubiquitin mutants (K11R/K48R and K27R/K11R/K48R). Incorporation of the K27R mutation into the K11R/K48R background suppressed the accumulation of ubiquitinated SLFN11 seen with K11R/K48R alone (**Fig. 5D**). Together, these findings suggest that K27-linked ubiquitylation acts upstream of, or in coordination with, other ubiquitin linkages to regulate SLFN11 (**Fig. 5E**), thereby influencing its chromatin recruitment and signaling functions in addition to its proteasomal degradation.

### RNF168 mediates SLFN11 ubiquitylation in SLFN11 middle/linker domain

We next sought to identify the E3 ligase responsible for SLFN11 ubiquitylation. Given that RNF168 is known to catalyze K27-linked ubiquitylation on histones H2A/H2AX during both unperturbed S phase and the DNA damage response, thereby promoting chromatin remodeling ^27,34^, we hypothesized that RNF168 may mediate SLFN11 ubiquitylation.

To test this, we transiently overexpressed Flag-tagged SLFN11 and performed immunoprecipitation using anti-Flag beads. RNF168 was readily detected in the SLFN11 pull-down, indicating a physical interaction (**Fig. 6A**). This association was validated by reciprocal immunoprecipitation using HA-tagged RNF168 as well as endogenous RNF168 and SLFN11 antibodies (**Fig. 6B, C**).

**Fig. 6.**
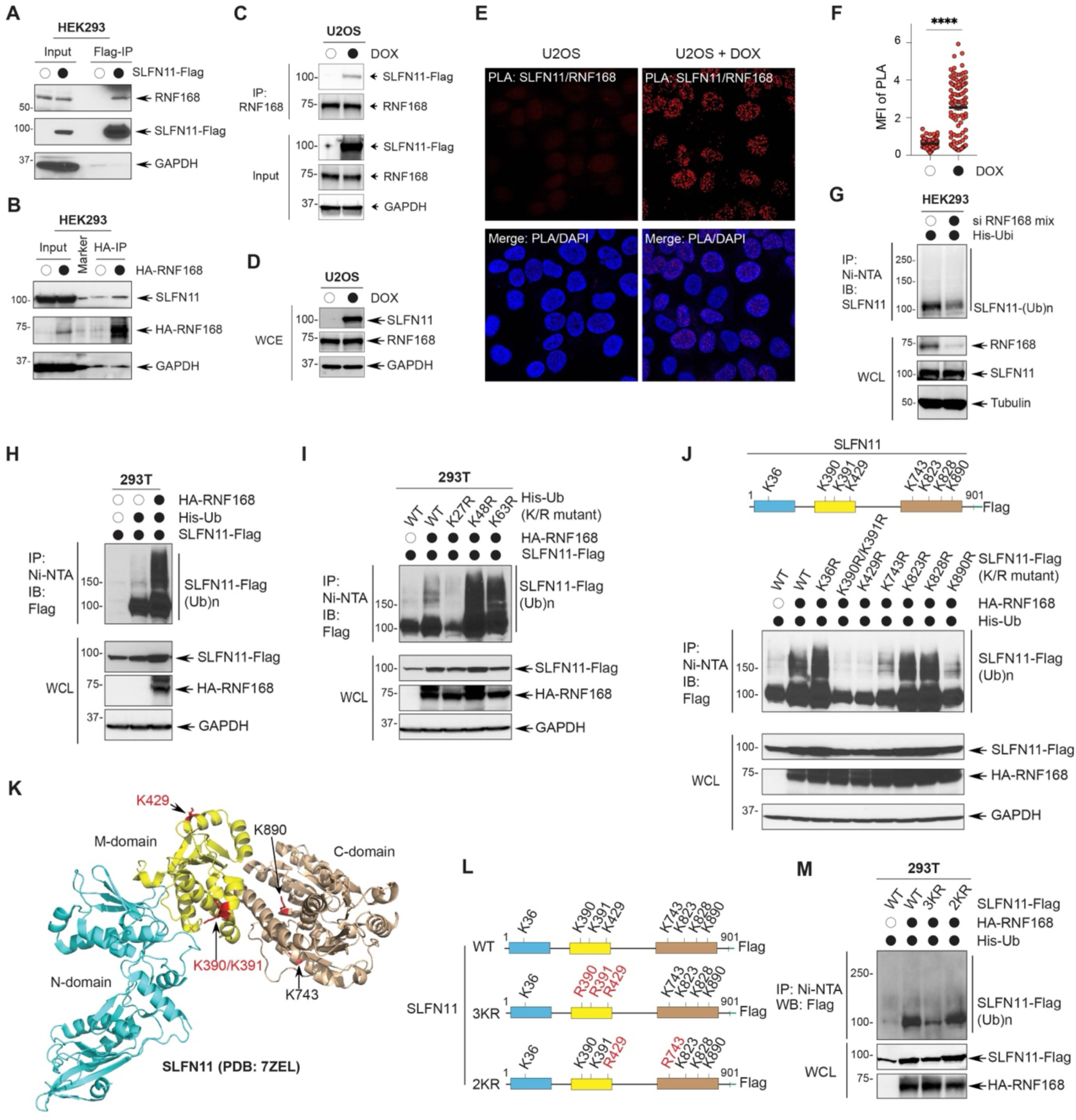
SLFN11 interacts with the E3 ubiquitin ligase RNF168 which targets the lysine residues in the middle linker domain of SLFN11. (A) Immunoprecipitation of RNF168 with SLFN11. HEK293 cells were transfected with Flag-tagged SLFN11 for 48 hrs. Cell lysates were immunoprecipitated with anti-flag M2 agarose beads. RNF168 was probed with anti-RNF168 antibody. (B) Immunoprecipitation of SLFN11 with RNF168. HEK293 cells were transfected with HA-tagged RNF168 for 48 hrs. Cell lysate were immunoprecipitated with anti-HA agarose beads. SLFN11 was probed with anti-SLFN11 antibody. (C) SLFN11 and RNF168 immunoblots of whole-cell lysate showing that SLFN11 expression does not affect RNF168 expression. DOX-inducible SLFN11-expressing U2OS cells were lysed after the DOX induction (72 hrs). (D) Immunoprecipitation of SLFN11 with an antibody against endogenous RNF168. After DOX induction for 72 hrs, U2OS cells were lysed and immunoprecipitated with anti-RNF168 with protein A/G agarose beads. SLFN11 was probed with anti-Flag M2 antibody. (E) Representative immunofluorescence images of proximity ligation assays (PLA) of SLFN11 and RNF168. DOX-inducible SLFN11-expressing U2OS cells were treated for 24 hrs with DOX and then incubated with primary antibodies against SLFN11 and RNF168. (F) Quantification of the PLA as in panel C. Points denote the mean fluorescence intensity (MFI) of PLA/cell with ± SEM (n > 80). ****p < 0.0001 (two-tailed unpaired t test). (G) RNF168-dependent ubiquitylation of SLFN11. HEK293 cells were transfected with His-Ub and siRNF168 mix. Cells were lysed and immunoprecipitated with Ni-NTA beads. Ubiquitylated SLFN11 [SLFN11-(Ub)n] was detected with anti-SLFN11 antibody. (H) Ubiquitylation of SLFN11 by RNF168. 293T cells were transfected with SLFN11-Flag, HA-RNF168, and His-Ub. Cells were lysed and immunoprecipitated with Ni-NTA beads. Ubiquitylated SLFN11 was detected with anti-Flag antibody. (I) K27-dependent ubiquitylation of SLFN11 by RNF168. 293T cells were transfected with SLFN11-Flag, HA-RNF168, and His-Ub (WT, K27R, K48R, and K63R). Cells were lysed and immunoprecipitated with Ni-NTA beads. Ubiquitylated SLFN11 was detected with anti-Flag antibody. (J) Mapping of the cellular ubiquitylation sites of SLFN11 by RNF168. Top: schematic representation of the tested lysine residues as potential targets for ubiquitylation by RNF-168 HEK293T cells were transfected with SLFN11-Flag, SLFN11-Flag K/R mutants, HA-RNF168, and His-Ub as indicated. Cells were lysed and immunoprecipitated with Ni-NTA beads. Ubiquitylated SLFN11 was probed with anti-Flag antibody. (K) Schematic representation of the location of the ubiquitylated lysine residues of SLFN11 (K390/K391, K429, K743, and K890) mapped onto SLFN11 protein structure (PDB: 7ZEL). (L) Schematic representation of the constructs used for the experiments shown in panel M. (M) Ubiquitylation sites of SLFN11 in its linker domain by RNF168. HEK293T cells were transfected with SLFN11-Flag, SLFN11-Flag K/R mutants (3KR:K390R/K391R/K429R, 2KR: K429R/K743R, as indicated), HA-RNF168, and His-Ub. Cells were lysed and immunoprecipitated with Ni-NTA beads. Ubiquitylated SLFN11 was probed with anti-Flag antibody.

We next examined whether SLFN11 and RNF168 interact in cells using proximity ligation assays (PLA) in doxycycline-inducible SLFN11-expressing U2OS cells. Nuclear PLA signals were observed upon SLFN11 induction, but not in non-induced controls (**Fig. 6E, F**), while RNF168 protein levels remained unchanged (**Fig. 6C**). These results indicate that SLFN11 and RNF168 interact within the nucleus.

Next, we determined whether RNF168 regulates SLFN11 ubiquitylation. Knockdown of RNF168 reduced SLFN11 ubiquitylation, whereas RNF168 overexpression enhanced it (**Fig. 6G, H**). This RNF168-associated increase in SLFN11 ubiquitylation was abolished by the K27R ubiquitin mutant (**Fig. 6I**). Conversely, a K27-only ubiquitin construct was sufficient to support SLFN11 ubiquitylation, unlike the K0 control (with all lysines mutated to alanine) (**Fig. S6**). From these results, we conclude that RNF168 mediates the SLFN11 ubiquitylation through K27-linkages.

To map the SLFN11 residues targeted by RNF168, we generated lysine-to-arginine mutants across potential SLFN11 ubiquitylation sites based on their location on SLFN11 surface. Mutations within the middle domain (K390R/K391R/K429R) and the C-terminal region (K743R/K890R) significantly impaired RNF168-driven ubiquitylation (**Fig. 6J**). Analyses of mutants with multiple lysine substitutions showed that the SLFN11-3KR triple mutant (K390R/K391R/K429R) exhibited a stronger reduction than the SLFN11-2KR mutant (K429R/K743R) (**Fig. 6K-M**). These results indicate that the middle/linker domain of SLFN11 is critical for RNF168-dependent modification. Collectively, these results identify RNF168 as a critical E3 ligase targeting SLFN11 for K27-linked ubiquitylation primarily within SLFN11’s middle/linker domain.

### RNF168-mediated ubiquitylation of SLFN11 is required for SLFN11 chromatin recruitment

We next tested whether RNF168-mediated ubiquitylation regulates SLFN11 chromatin recruitment. In response to VLX-1570, we observed coordinated chromatin accumulation of RNF168 together with SLFN11, with VLX-1570 inducing a more pronounced enrichment of both RNF168 and SLFN11 compared to CPT (**Fig. 7A-C**), suggesting that RNF168 is actively engaged at sites where SLFN11 is recruited. Consistent with this, siRNA-mediated depletion of RNF168 significantly impaired SLFN11 chromatin recruitment under both VLX-1570- and CPT-induced conditions (**Fig. 7A-C and S7**), suggesting that RNF168-dependent ubiquitylation as a key upstream event required for efficient SLFN11 loading onto chromatin in response to DUB inhibition as well as DNA damage.

**Fig. 7.**
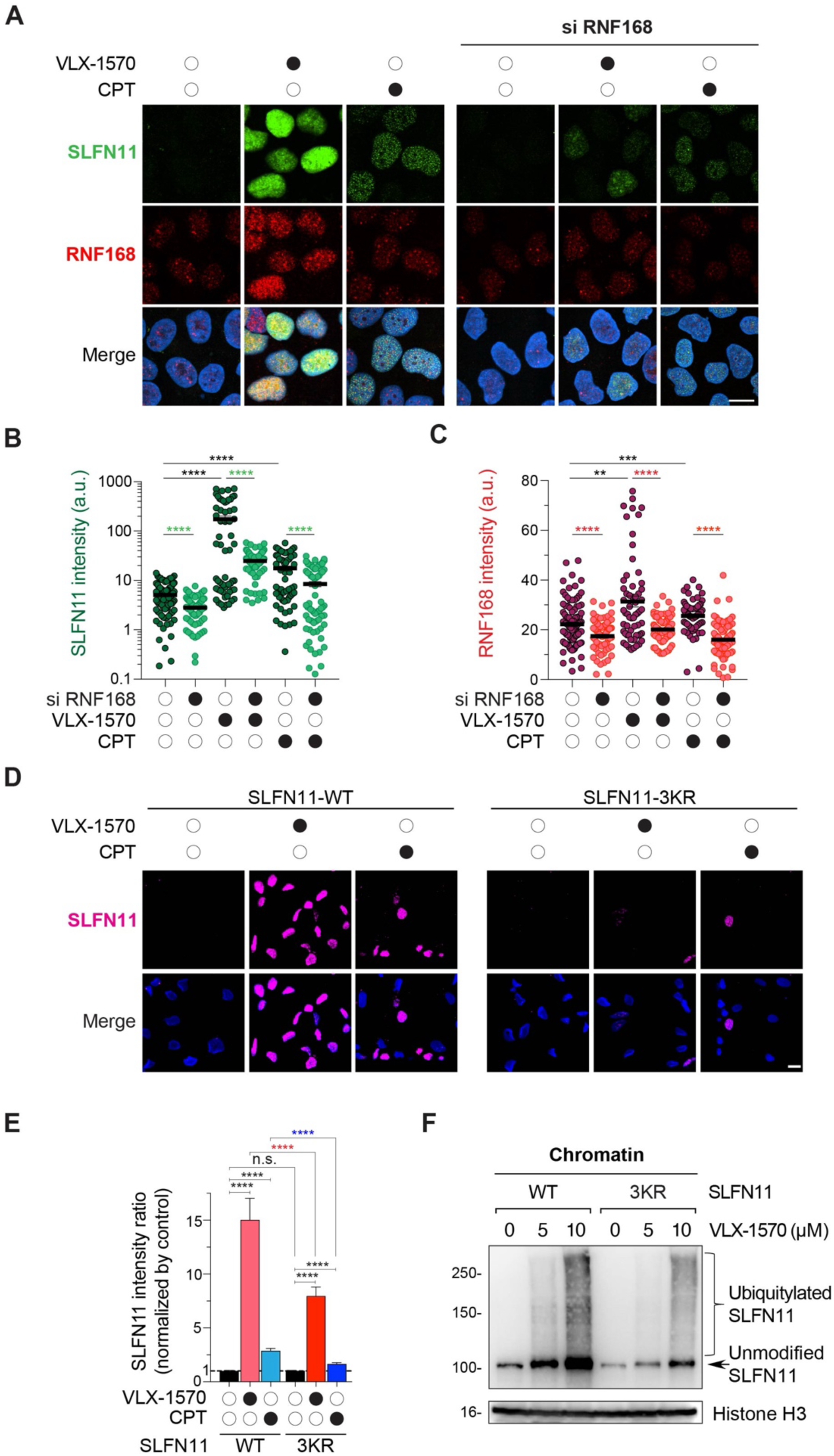
RNF168 and the ubiquitylation sites of SLFN11 in its middle domain promotes chromatin recruitment of SLFN11. (A) Representative immunofluorescence images showing chromatin-bound SLFN11 (green), RNF168 (red) and DAPI (blue). DOX-inducible SLFN11-expressing U2OS cells were treated with VLX-1570 (1 µM) or CPT (10 µM) for 2 hrs after DOX induction (72 hrs) and siRNF168 transfection (48 hrs). Scale bar: 5 µm. (B, C) Quantification of chromatin-bound SLFN11 and RNF168 signals in individual cells for the indicated treatments (n = 57 – 97 cells per condition). **p < 0.01. ***p < 0.001. ****p < 0.0001 (two-tailed unpaired t test). a.u., arbitrary units. (D) Representative immunofluorescence images showing chromatin-bound SLFN11 (magenta) and DAPI (blue) in DOX-inducible SLFN11 clones (WT, 3KR) of U2OS cells. Cells were treated with VLX-1570 (1 µM) or CPT (10 µM) for 2 hrs. Scale bar: 10 µm. (E) Quantification of chromatin-bound SLFN11 fluorescence (average per condition) for the treatment depicted in panel D (mean ± SEM, N = 411 – 690 cells per condition). ***p < 0.0001 (two-tailed unpaired t test). (F) SLFN11 immunoblots of chromatin fraction. DOX-inducible SLFN11 clones (WT, 3KR) of U2OS cells were treated with VLX-1570 (5 µM, 10 µM, 2 hrs) after the DOX induction (72 hrs), lysed and fractionated with pre-extraction buffer.

To test whether the importance of SLFN11 ubiquitylation, we used our doxycycline-inducible U2OS cells expressing a ubiquitylation-deficient SLFN11 mutant (3KR: K390R/K391R/K429R). Chromatin recruitment of SLFN11 upon VLX-1570 or CPT treatment was suppressed in cells expressing the 3KR mutant compared to wild-type SLFN11 (**Fig. 7D, E**). Consistently, SLFN11 ubiquitylation was diminished in 3KR mutant cells (**Fig. 7F**) despite comparable SLFN11 expression between wild-type and mutant conditions (**Fig. S7B-C**). Collectively, these results establish that RNF168-mediated ubiquitylation of SLFN11, particularly within its middle/linker domain, is critical for SLFN11 chromatin recruitment.

## DISCUSSION

SLFN11 was previously known to be recruited to stressed replication forks within a few hours of DNA damage induced by chemotherapeutic agents, where it enforces a global and irreversible replication block that ultimately results in cell death ^13–16,20,35,36^. In the present study, we expand this paradigm by identifying an additional mechanism recruiting SLFN11 to chromatin. Using a high-throughput imaging (HTI) screening strategy with small-molecule inhibitors targeting defined signaling and cancer-relevant pathways, we found that deubiquitinase (DUB) inhibitors are potent inducers of SLFN11 chromatin association. Mechanistically, we demonstrate that RNF168-mediated ubiquitylation within SLFN11’s middle linker domain is essential for the recruitment of SLFN11 to chromatin.

Deubiquitylation, which is catalyzed by ∼100 enzymes across approximately six major DUB families, plays a central role in protein homeostasis and signaling ^37^. Beyond protein turnover, ubiquitin signaling regulates chromatin architecture and coordinates the recruitment of DNA repair and chromatin-associated regulatory factors ^38–43^. Key chromatin mediators, through ubiquitylation, can integrate into target sites as protein-DNA complexes or multi-protein complexes, forming molecular scaffolds ^25^ and enzyme complexes ^44^. Our screening with drugs targeting the genome and epigenome identified multiple DUB inhibitors—including VLX-1570, PR-619, and WP1130—as the strongest inducers of SLFN11 chromatin recruitment in comparison of the established inducers of SLFN11 nuclear foci included in the screen (doxorubicin, CPT, the Wee1 inhibitor AZD-7762 or the CHK1 inhibitor Prexasertib). This observation highlights the importance of ubiquitin dynamics in controlling SLFN11 localization. We also show that DUB inhibitors promotes SLFN11 accumulation at open, transcriptionally active chromatin regions enriched for H3K27 hyperacetylation, and functionally suppresses RNA synthesis and that inhibition of ubiquitin activation using TAK-243 abolishes SLFN11 chromatin recruitment.

The chromatin recruitment of SLFN11 induced by DUB inhibitors differs from its recruitment in response to replication stress. Whereas replication stress drives SLFN11 into discrete nuclear foci that colocalize with RPA-coated single-stranded DNA ^13,14^, DUB inhibitors induces a broad, pan-nuclear chromatin association that does not colocalize with RPA and is not associated with detectable γH2AX formation. These findings suggest that SLFN11 recruitment to chromatin is not restricted to sites of stalled replication forks but can also be globally redistributed across remodeled chromatin landscapes. This expanded mode of recruitment implies broader roles for SLFN11 in genome regulation, potentially through interactions with chromatin-associated factors that influence DNA replication ^16^, chromatin accessibility ^18^, and transcriptional control ^36,45^. Accordingly, ChIP–seq profiling revealed that SLFN11 predominantly occupies open, transcriptionally active chromatin regions. Therefore, SLFN11 appears to act as a dual-function protein, playing a surveillant role in cellular processes such as normal transcription and ribosome biogenesis in an less RPA-dependent context ^36,46^, while also responding to DNA damage in an RPA- and ssDNA-dependent manner.

Mechanistically, we identify K27-linked polyubiquitylation as the regulatory signal controlling SLFN11 chromatin recruitment and turnover. While K48- and K63-linked ubiquitin chains are classically associated with proteasomal degradation^25^, our data suggest that K27-linked ubiquitination also acts as an upstream regulatory mark promoting SLFN11 chromatin association and subsequently facilitating its modification by K48/K63 linkages for proteasomal turnover. This is consistent with emerging roles of K27-linked ubiquitin chains in chromatin remodeling and DNA damage responses ^25,27,41^.

Given the established preference of RNF168 for generating K27-linked ubiquitin chains and its well-characterized functions in chromatin-associated DNA replication and repair processes^34,41^, we investigated its role in SLFN11 regulation. Our findings demonstrate that RNF168 is a critical E3 ligase for SLFN11 ubiquitylation, required for its chromatin recruitment upon DUB inhibition. We mapped the ubiquitylation sites to lysine residues K390, K391, and K429 within the previously uncharacterized linker domain between the N-terminal RNAse and C-terminal DNA binding domains of SLFN11. While the N-terminal nuclease domain and C-terminal ATPase/ssDNA-binding domains have been extensively studied^1,41^, our results highlight the linker region as a previously unrecognized regulatory region for ubiquitin-dependent chromatin engagement. This mechanism may involve cooperation with chromatin-associated partners such as the DNA/RNA helicase DHX9 ^47^, which has been implicated in RNF168-dependent ubiquitin signaling ^41,48^, suggesting that the importance of SLFN11 ubiquitylation in driving SLFN11’s interactions with chromatin cofactors, drawing parallels with SLFN12-PDE3A regulation ^49,50^.

From a translational perspective, our findings suggest that SLFN11 is a potential functional mediator of DUB inhibitor activity in cancer therapy. Although several DUB inhibitors have entered clinical evaluation, including the USP1 inhibitor KSQ-4279 (NCT05240898) ^51^, and the dual UCHL5/USP14 inhibitor VLX-1570 (NCT02372240) ^52^, clinical development has been limited by safety and toxicity concerns, including dose-limiting adverse effects. These observations underscore the need for improved therapeutic strategies and a deeper understanding of SLFN11-associated ubiquitin signaling pathways that may influence drug response and selectivity.

In summary, we identify a ubiquitin-dependent mechanism governing SLFN11 chromatin recruitment upon DUB inhibition, requiring RNF168-mediated K27-linked ubiquitylation. This pathway drives the recruitment of SLFN11 to open chromatin, expanding its functional role beyond replication stress. Future work should define the upstream DUBs and assess the therapeutic relevance of this regulatory axis in cancer.

## MATERIALS AND METHODS

### Automated image acquisition and high throughput image analysis

A Yokogawa CV7000S high-throughput spinning-disk confocal microscope was employed for automated imaging of stained 384-well plates. DAPI and chromatin-bound SLFN11 were detected using 405 nm and 561 nm excitation lasers, respectively. Imaging in both channels utilized a 405/488/561/640 nm excitation dichroic mirror and a 40× air objective lens (NA 0.95). Sequential single-plane images were captured using a 16-bit sCMOS camera (2560×2160 pixels, 2×2 binning, pixel size 0.325 μm) with bandpass filters (445/45 nm for DAPI, 525/50 nm for SLFN11). Proprietary Yokogawa software was employed for background and illumination correction, and images were saved in TIFF format. For high-throughput image analysis, image files were processed using Columbus 2.7.1 software (PerkinElmer). The analysis pipeline segmented nuclei using DAPI staining and quantified SLFN11 fluorescence intensity within nuclear regions of interest (ROIs) after treatment with 162 compounds. Nuclei with roundness below 0.775 (typically segmentation artifacts) and those touching the image border were excluded. The mean SLFN11 nuclear fluorescence intensity was averaged across all cells per well, and well-level results were exported as .txt files for downstream analysis. This platform was used in Fig. 1, S1A, 7D, and 7E.

### Pre-extraction for the detection of chromatin-bound SLFN11

For biochemical cell fractionation, the cells were incubated with 40 µL of 0.1% Triton X-100 CSK buffers (10 mM PIPES [pH 6.8], 100 mM NaCl, 300 mM sucrose, MgCl2, 1 mM EGTA, 1 mM EDTA, 1 mM, and 0.1% Triton X-100) per well in a 6-well plate for 5 minutes on ice 53. The samples were then centrifuged at 1500 g for 5 minutes, followed by a single PBS wash at 1500 g for 5 minutes. The supernatant, containing nuclear and cytoplasmic soluble proteins, was collected. The pellet (chromatin fraction) was sequentially resuspended in 20 µL PBS and then in 20 µL of 2X boiling lysis buffer (50 mM Tris-Cl [pH 6.8], 2% SDS, and 850 mM β-mercaptoethanol). The mixture was boiled for 10 minutes. Post-boiling, 20 µL of the sample was mixed with 20 µL of 2X SDS sample buffer and boiled for an additional 5 minutes. The protein samples were then resolved on an 8% SDS-PAGE gel for subsequent Western blot analysis.

### Cell lines

HEK293 (CRL-1573), 293T (CRL-3216), DU145 (HTB-81) and U2OS (HTB-96) cell lines were purchased from American Type Culture Collection (ATCC) and maintained without further authentication. All cell lines were grown in DMEM medium (Thermo Fisher Scientific) containing 10% FBS and 1% penicillin-streptomycin. U2OS doxycycline-inducible SLFN11 expressing cells were established by virus infection containing pLVX-TetOne-blasticidin-3X Flag-SLFN11. After 48 hours post-infection, cells were selected with blasticidin (10 µg/ml) (Gibco). For experiments, cells were prepared at low passages within 15 passages and were routinely tested for mycoplasma contamination (MycoAlert Mycoplasma Detection Kit_Lonza).

### Reagents

162 compounds used in the screening, as well as other DUB inhibitors used in validation in Fig. S1B and S1C were provided by Division of Preclinical Innovation (NCI, NIH). MG132 (C2211) was purchased from Millipore-Sigma. Camptothecin was acquired from the Developmental Therapeutics Program (DCTD, NCI, NIH). TAK-243 (#HY-100487) was purchased from MedChemExpress.

### Immunofluorescence staining analysis with or without EdU Labeling

Cells were incubated with 10 mM 5-ethynel-20-deoxyuridine (EdU) for 30 min just before collection if needed. Cells (5×104 cells) were seeded on 4 or 8-well chamber slides (Nunc™ Lab-Tek™ II CC2™, Thermo Fisher, #154917, #154941). Cells were treated with drugs as described in the figure legends. Cells were then pre-extracted in 0.1% Triton X-100 cytoskeleton (CSK) buffer (10 mM Tris [pH 6.8], 100 mM NaCl, 300 mM sucrose, MgCl2, 1 mM EGTA, 1 mM EDTA, and 0.1% Triton X-100) for 5 min on ice, following fixation in 4% PFA for 15 min. After a subsequent blocking step in 5% BSA/PBS for 1h, cells were incubated with the following primary antibodies overnight incubation at 4 DC (dilution 1:1000): anti-SLFN11 (Santa Cruz Biotechnology, D2) or anti-SLFN11 (Cell Signaling, D8W1B), anti-H3K27ac (Cell Signaling, D5E4), anti-H3K27me3 (Cell Signaling, C36B11), anti-Ub (Santa Cruz Biotechnology, P4D1), anti-FK2 (Millipore Sigma, 04-263), anti-RPA2 (Santa Cruz Biotechnology, 9H8), anti-γH2AX (Millipore Sigma, 05-636), and anti-RNF168 (Millipore Sigma, ABE367) in 5% BSA/PBS. Next day, cells were washed three times with 1X PBS and incubated for 1h with either anti-mouse (Invitrogen, A-11004) or anti-rabbit (Invitrogen, A-11034) alexa fluor secondary antibodies in 5% BSA/PBST (dilution 1:1000) for 1 h. After three times washing with 1X PBS, cells were mounted with VECTASHIELD® Antifade Mounting Medium with DAPI (H-1200; Vector laboratories). For EdU analysis, Click-iT EdU (5-ethynyl-2′-deoxyuridine) Alexa Fluor 488 Imaging Kit (Thermo Fisher Scientific) were performed according to the manufacturer’s instructions.

Except for Fig. 1D, S1A, 6E, and 7D, fluorescence images were captured by using Nikon SoRa spinning disk confocal microscope with 60 objective lens. For Fig. 6E, fluorescence images were captured by Zeiss LSM 880 confocal microscope with 63x objective lens.

### Immunofluorescence Microscopy Image Analysis

Except for Fig. 1, S1A, 7D, and 7E, signal intensities within individual cells were quantified using ImageJ. A standardized circular region detected by DAPI staining was applied consistently across all isogenic cell line samples to measure mean signal intensities. These values were used to generate individual signal and co-localization plots.

Signal distribution profiles are shown in Fig. 4B. The resulting data were then exported to GraphPad Prism 10 (GraphPad Software) for visualization. A signal intensity threshold of 3 was applied to both EdU and SLFN11 channels to eliminate background noise in Fig. S3E.

### EU labeling and quantification in immunofluorescence assays

Following drug treatment as described in the figure legends, cells were incubated with 5-ethynyl uridine (EU) for 1 hour at 1 mM concentration. Nascent RNA synthesis was detected using the Click-iT RNA Alexa Fluor 488 Imaging Kit (Thermo Fisher, C10329) according to the manufacturer’s instructions. EU fluorescence intensities were quantified using Fiji, and data were visualized using GraphPad Prism 10.

### Plasmids and site-directed mutagenesis

pCI-His-hUbiquitin (a gift from Astar Winoto (Addgene, #31815), pRK5-HA-Ubiquitin K0 (7A) (Addgene, #22902), and pRK5-HA-Ubiquitin 27 (6A) (Addgene, #17603) were purchased from Addgene and ubiquitin K/R mutants were generated by using site-directed mutagenesis with the primers (Table S2). pcDNA3-SLFN11-Flag was generated by amplifying SLFN11 cDNA (HsCD00082389) in the pDONR201 vector (DNASU plasmid repository) together with primers (Table S2) including HindIII and ApaI restriction enzyme sites. SLFN11 K/R mutants were generated by using site-directed mutagenesis with the primers (Table S2). pcDNA3-HA-RNF168 was generated by amplifying RNF168 cDNA (HsCD00334902) in the pCMV-SPORT6 vector (DNASU plasmid repository) together with primers (Table S2) including BamHI/BglII and XhoI restriction enzyme sites.

### Chromatin immunoprecipitation sequencing (ChIP-seq) and data analysis

Chromatin was crosslinked with formaldehyde (final concentration 1%) for 15 min at room temperature and quenched with glycine (final concentration 0.125 M). Nuclei were isolated using Farnham lysis buffer (5 mM PIPES, pH 8.0; 85 mM KCl; 0.5% NP-40) supplemented with PMSF and protease inhibitor cocktails. Chromatin was sheared to an average fragment size of 200–500 bp using a Misonix Sonicator 3000 (30 cycles of 20 s on/30 s off).

Sonicated chromatin was immunoprecipitated with Dynabeads Protein A (Invitrogen, #10002D) conjugated to the indicated antibodies. Antibodies used for ChIP-seq included anti-Flag M2 (MilliporeSigma, #F1804) and anti-H3K27ac (Active Motif, #39133). Following sequential bead washes, immunoprecipitated chromatin was reverse crosslinked overnight at 65 °C in the presence of 1% SDS and 1 mg/mL proteinase K. DNA was purified using the QIAquick PCR Purification Kit (Qiagen, #28004).

Purified DNA fragments were end-repaired and ligated to Illumina adapters using the NEBNext Ultra II DNA Library Prep Kit for Illumina (New England BioLabs, #E7103L and #E6440). Libraries were quantified using a TapeStation system (Agilent Technologies) and sequenced on a NextSeq 2000 platform with 101-bp single-end reads. Base calling and demultiplexing were performed using Illumina DRAGEN BCL Convert software (v4.2.7).

The raw single-end ChIP-seq reads were trimmed for adapters using Trimmomatic (v0.39) software and then mapped to reference human genome (version hg38) using BWA (v 0.7.17) 54. PCR duplicates were removed using Picard (v3.4.0) and reads aligned to ENCODE hg38 blacklist were filtered out. High-confidence peaks were identified by MACS2 (v2.2.7.1)55 with the narrow algorithm at p-value ≤ 1×10⁻⁵. Peak sets were annotated against hg38 using ChIPseeker (v 1.38.0) with the promoter defined as transcription starting site +200/− 2000 bp. Genome-wide coverage tracks were generated from blacklist-filtered BAMs using bamCoverage function from deepTools (3.5.6)56 with 25-bp binning and RPKM normalization, producing bigWig files. These bigwig files were also used to plot heatmaps using deepTools (v3.5.6).

The ChIP-seq data generated in this study have been deposited in the Gene Expression Omnibus (GEO) database under accession code GSE318424.

### Proximity ligation assay

Doxycycline (Dox)-inducible SLFN11 expressing U2OS cells were seeded in 4-well Nunc™ Lab-Tek™ II CC2™ Chamber Slide System (154917; Thermo Fisher) and grown for 24 hours with Dox 10 μg/ml. After cell fixation, cells were permeabilized in 0.3% Triton X-100/1xPBS and analyzed with Duolink™ In Situ Red Starter Kit Mouse/Rabbit (DUO92101, Sigma Aldrich) according to the manufacturer’s instructions. Briefly, cells were blocked and incubated with anti-SLFN11 antibody and rabbit anti-RNF168 antibody for 1 hour at room temperature. Cells were labeled with PLUS and MINUS PLA probes and subsequently ligated and amplified. Finally, cells were mounted with the Duolink In-Situ Mounting Medium with DAPI. PLA and DAPI signals were captured by Zeiss LSM 780 ELYRA (63x magnification) microscopy and quantified by ImageJ software.

### Small interfering RNA (siRNA) transfection

siRNAs targeting RNF168 were purchased from Qiagen. ON-TARGETplus Human USP14 and UCHL5 siRNA SMARTpools (L-006065-00-005 for USP14, M-006060-03-0005 for UCHL5; Dharmacon/Horizon Discovery) were transfected using Lipofectamine RNAiMAX (Invitrogen) according to manufacturer’s instructions. The ON-TARGETplus Human USP14 and UCHL5 siRNA SMARTpools each consists of four individual siRNAs targeting distinct regions of the respective mRNAs. The siRNAs were pooled at equimolar concentrations and used for all knockdown experiments. The non-targeting siRNA (siControl) was obtained from Origene and used as the control. The siRNA sequences are listed in Table S3.

### In vivo ubiquitylation assay

HEK293 and 293T cells were transfected with His-conjugated ubiquitin plasmids and the plasmids indicated in figure legends for 48 hours with/without MG132 (20 µM) and lysed by the denatured buffer A (6 M guanidine-HCl, 0.1 M Na2HPO4/NaH2PO4, 10 mM imidazole). The cell extracts were separated into the whole cell lysate controls and immunoprecipitation samples, which were then incubated with Ni-NTA (Qiagen) beads for 3 hours. The beads were washed once with buffer A, twice with buffer A/TI (Ratio 1:3), and twice with TI (25 mM Tris-HCI, pH 6.8, and 20 mM imidazole), and finally subjected to immunoblotting analysis with the indicated antibodies.

### Immunoprecipitation

Cells were lysed with NETN100 buffer (1% NP40, 100 mM NaCl, 0.1 mM EDTA, and 50 mM Tris [pH 7.5]) supplemented with 1X protease inhibitor cocktail and 1 μl benzonase (EMD Millipore) for 40 min on ice. Cells were then centrifuged at 15,000 rpm for 10 min. Cell lysates were separated into input controls (5%) and immunoprecipitation samples that were incubated with anti-Flag M2 affinity gel (Sigma, #A2220), anti-HA agarose (Sigma, #A2095), and Protein A/G Plus Agarose (Thermo Scientific, #20423) in NETN100 buffer overnight at 4°C. After then, the immune complexes were washed three times with NETN100 buffer.

### Western blotting analysis

Cells were lysed with NETN300 buffer (1% NP40, 300 mM NaCl, 0.1 mM EDTA, and 50 mM Tris [pH 7.5]) supplemented with protease inhibitor cocktail for 40 min on ice. After centrifugation for 10 min at 15,000 rpm, cell lysates were resolved by SDS-PAGE gels and transferred onto PVDF membranes (Millipore). Membranes were then blocked in 5% milk/PBS-T and immunoblotted with the following antibodies: anti-SLFN11 (dilution 1:1000, D2, Santa Cruz Biotechnology, sc-51507), anti-USP14 (dilution 1:1000, Thermo Fisher, A300919A), anti-UCHL5 (dilution 1:1000, Thermo Fisher, PA5-110544), anti-Ub (dilution 1:1000, Santa Cruz Biotechnology, P4D1), anti-RNF168 (dilution 1:1000, Millipore Sigma, 06-1130-I), anti-⍺-Tubulin (dilution 1:1000, Cell Signaling, #2125), anti-GAPDH (dilution 1:1000, #2118), anti-Histon H3 (dilution 1:5000, Cell Signaling, D1H2), anti-HA tag (dilution 1:1000, #3724), anti-Flag (dilution 1:1000, #F1804).

After overnight incubation at 4°C, membranes were incubated with a peroxidase (HRP)-conjugated sheep anti-mouse antibody (dilution 1:4000, GE Healthcare, NA9310) and anti-rabbit antibody (dilution 1:4000, GE Healthcare, NA9340) for 1 h at room temperature. Finally, membranes were washed three times with PBS-T buffer and were developed by using SuperSignalTM West Chemiluminescent Substrates (Thermo Scientific). Chemiluminescent detection was performed by ChemiDoc MP Imaging System (Bio-rad).

### Statistical analysis

All statistical analyses were conducted with GraphPad Prism 10. Test methods are described in each figure legend.

## RESOURCE AVAILABILITY

### Lead contact

Requests for further information and resources should be directed to and will be fulfilled by the lead contact, Yves Pommier.

### Materials availability

All materials, reagents, and detailed protocols generated in this article are available upon request to the lead contact.

### Data and code availability

This paper does not report original code. Any additional information required to reanalyze the data reported in this paper is available from the lead contact upon request.

## ACKNOWLEDGMENTS

This work was supported by National Cancer Institute Center for Cancer Research intramural grants Z01 BC 006161 and Z01 BC 006150 (to Y.P.). We thank Dr. Anagh Ray (DTB/CCR/NCI/NIH), Dr. Hye Kyung Lee (NIDDK/NIH) and Dr. Madeline Wong (CCR Genomic Core/NCI/NIH) for ChIP-seq expertise and discussion.

## AUTHOR CONTRIBUTIONS

Conceptualization, D.T., U.J., Y.P.; methodology, D.T., U.J., G.P., L.O., A.D.T.; investigation, D.T., U.J., S.N.H., L.K.S.; resources, C.J.T.; formal analysis, D.T., U.J., Y.W., G.P.; visualization, G.P., L.O., Y.W.; supervision, U.J., Y.P.; writing– original draft, D.T., U.J., Y.P.; and writing– review & editing, D.T., J.M., U.J., and Y.P.

## DECLARATION OF INTERESTS

The authors declare no competing interests.

